# Design And Optimization Studies Of Tablet In Tablet Formulation Of Diclofenac And Misoprostol: Application Of Response Surface Methodology And Compressional Behaviour Strategy

**DOI:** 10.1101/2022.11.04.515246

**Authors:** Shabana Naz Shah, Huma Ali, Riffat Yasmin, Shaheen Perveen, Farya Zafar, Fozia Israr, Nousheen Aslam

**Affiliations:** Department of Pharmaceutical Chemistry, Faculty of Pharmacy SBB Dewan University, Karachi, Pakistan; Department of Pharmaceutics, Institute of Pharmaceutical Sciences, Jinnah Sindh Medical University, Karachi, Pakistan; Dow College of Pharmacy, Dow University of Health Sciences, Karachi, Pakistan; Department of Pharmaceutics, Faculty of Pharmacy, University of Karachi, Karachi, Pakistan; Mohammed Al-Mana College for Medical Sciences, Dammam, Saudi Arabia

**Keywords:** Compression coated tablets, Diclofenac sodium, Misoprostol, Enteric coated

## Abstract

The present study was aimed to develop compression coated tablets of diclofenac sodium (75mg) in the inner core and misoprostol (200 mg) as the outer shell for the effective treatment of rheumatoid arthritis as this dosing frequency is not available in the market yet. Diclofenac sodium inner tablet was manufactured by wet granulation and its enteric coating was applied by Eudragit L 100-55, Isopropyl Alcohol and Opadry II Blue 85 F205034. While immediate-release misoprostol outer shell was also manufactured by wet granulation and coated by Opadry white and Polyethylene glycol 6000. Design of experiment^®^ software was used for formulation design and optimization. Quality attributes such as tablet weight, hardness, disintegration time, percent drug dissolution and assay were performed as per official methods and satisfactory results were reported. Physical and chemical stability of selected formulations was evaluated following the ICH guidelines for accelerated stability testing. The compressional analysis of optimized formulation was performed to check the optimum compression pressure to obtain a stable formulation. Based on satisfactory quality attributes; formulation DF9MF7 was successfully developed as compression coated tablet with calculated shelf life of 4.8years. Compression coated tablets comprising of enteric coated diclofenac sodium as inner core and misoprostol as outer shell were successfully developed by wet granulation.

## 1. Introduction

Tablet in tablet technology or compression coated tablet was introduced as an alternative to coating technique [1]. Compression coated tablet contains two layers where the initial layer is formed by a slight compression of the drug along with the powder known as the internal core; while granules and powder of the second coat are then transferred into the tableting machine containing initially compressed tablet to form a constant mass around surfaces and edges of the initially compressed tablet known as outer shell resulting in the finished product [2]. The outer shell serves to provide tablet stability and it controls the drug release [3]. The major advantage of tablet in tablet formulation included the separation of incompatible material in the core and outer shell. The pharmacokinetic interaction between the two concomitantly administered medications can be avoided by formulating tablet in tablet dosage form with the time interval in their release using different polymeric excipients. This is of particular significance when the drug has a bitter taste or it is unstable under acidic conditions or causes irritation to the stomach [1,4,5]. Despite the advantages offered by a tablet in tablet technology, there are several challenges that include the possibility of cross-contamination between the layers. There is often a possibility of elastic modulus mismatch between the adjacent layers leading to the inadequate attachment of the second layer.

Diclofenac sodium pertains to the class of non-steroidal anti-inflammatory drugs (NSAIDs) that works through the inhibition of COX-1 and COX-2 enzymes. It is used for the management of osteoarthritis, ankylosing spondylitis, and rheumatoid arthritis due to its potent analgesic activity. The half-life of diclofenac sodium is short i.e. 2 hours [6]. However Diclofenac sodium causes gastrointestinal adverse events such as inflammation, bleeding, ulceration and perforation of intestinal wall [7,8]. Therefore enteric coating films are widely used to protect the stomach from such effect through providing a surface that remain unchanged and exceedingly stable in stomach having acidic environment but releases drug readily in intestine containing basic environment [9]. These films are prepared by using polymers which are soluble in pH value more than 5-6. The chiefly utilized and recommended polymers used for pH-sensitive enteric coating are hydroxypropyl methyl cellulose phthalate, methacrylic acid copolymers, cellulose acetate phthalate, cellulose acetate succinate, polyvinyl acetate phthalate, hydroxylpropyl ethyl cellulose phthalate, cellulose acetate tetrahydrophthalate, and acrylic copolymers [10-12]. Eudragit is preferred choice for enteric coating because of their nontoxic and nonirritant nature that offers versality. Furthermore, the process of coating with eudragit is safer, easier and economical.

Misoprostol is an orally active analogue of prostaglandin E1 that possess antisecretory and cytoprotective property [13]. Misoprostol is recommended by FDA for the management of NSAID-induced gastric and duodenal ulcers.[14]. The misoprostol formulation is difficult to formulate due to its viscous liquid form and instable nature at room temperature. Misoprostol undergoes dehydration process under acidic or alkaline condition containing small quantity of water resulting in elimination of 11-hydroxy group. This results in formation of an A-type prostaglandin that can further isomerize to form prostaglandin B-form. Misoprostol can be stabilized by the use of solid dispersion of misoprostol in HPMC resulting in decreased mobility of water [15,16]. The literature review indicated that diclofenac sodium and misoprostol showed comparable absorption rate, no sign of interaction and do not accumulate in plasma after recommended dose [17]. Therefore, the present study was aimed to develop diclofenac sodium in the inner core and misoprostol as the outer shell for the effective treatment of rheumatoid arthritis.

## 2. Materials and methods

### 2.1 Materials

Diclofenac sodium was received as a gift from Sami Pharmaceutical (Pvt.) Ltd. Lactose monohydrate and Sodium Starch Glycolate, Povidone K-30, Corn Starch, Magnesium Stearate, and Isopropyl Alcohol were procured from FMC Corp, USA and Merck, Darmstadt, Germany. For enteric coating of the core tablets Eudragit L 100-55 (Evonik Industries, GmbH, Germany), Opadry II Blue 85 F205034 (Colorcon, Verna, Goa, India), Isopropyl Alcohol (Merck, Darmstadt, Germany), and Castor oil were procured form Colorcon. Misoprostol was purchased from Atco Pharmaceuticals. Micro crystalline cellulose (102), Aerosil-200, Magnesium Stearate and Talc were purchased from FMC Corp, USA and Merck, Darmstadt, Germany.

### 2.2 Equipments

Top Loading Balance (Snow Rex, Taiwan), Tablet Compression Machine (Express 30), homogenizer (Daihan Sci, USA), Planetary Mixer (Kenwood, UK), Hot Air Oven (Wiseven, Germany), Digital Vernier Calliper (Karl Kolb, Germany) Disintegration Tester (Galvano Sci, LHR, Pak), Friabilator (Copley, UK), Hardness Tester (Galvano Sci, LHR, Pak), Dissolution Tester (Multi Paddle) (Copley, UK), pH meter (WTW, Germany), UV Spectrophotometer UV-1800 (Schimadzu, Japan), Coating Pan-CP-9 with spray gun (Karl Kolb, Pharma G, Germany), Binary Gradient High Performance Liquid Chromatography (LC-20 Gradient, Schimadzu, Japan). Pump (LC-10AS, Shimadzu), UV detector (SPDA-10A, Shimadzu) and Lichrospher100 column (250×4.0 mm, RP-8, 5μm).

### 2.3 Methods

#### 2.3.1 Micromeritic Evaluation

Granules of Diclofenac sodium and Misoprostol were separately manufactured by wet granulation and then subjected to micromeritic examination such as tapped density, bulk density, angle of repose, compressibility index and Hausner’s ratio as mentioned in USP[18].

#### 2.3.2 FT-IR Studies

FT-IR spectra of pure drugs and optimized formulation blends were analyzed in the region of 500 - 4000 cm^-1^. Powder blends in 1:1 ratio of API and excipients were placed directly as a thin film for analysis using FT-IR Spectrophotometer (Figure 1).

**Figure 1:**
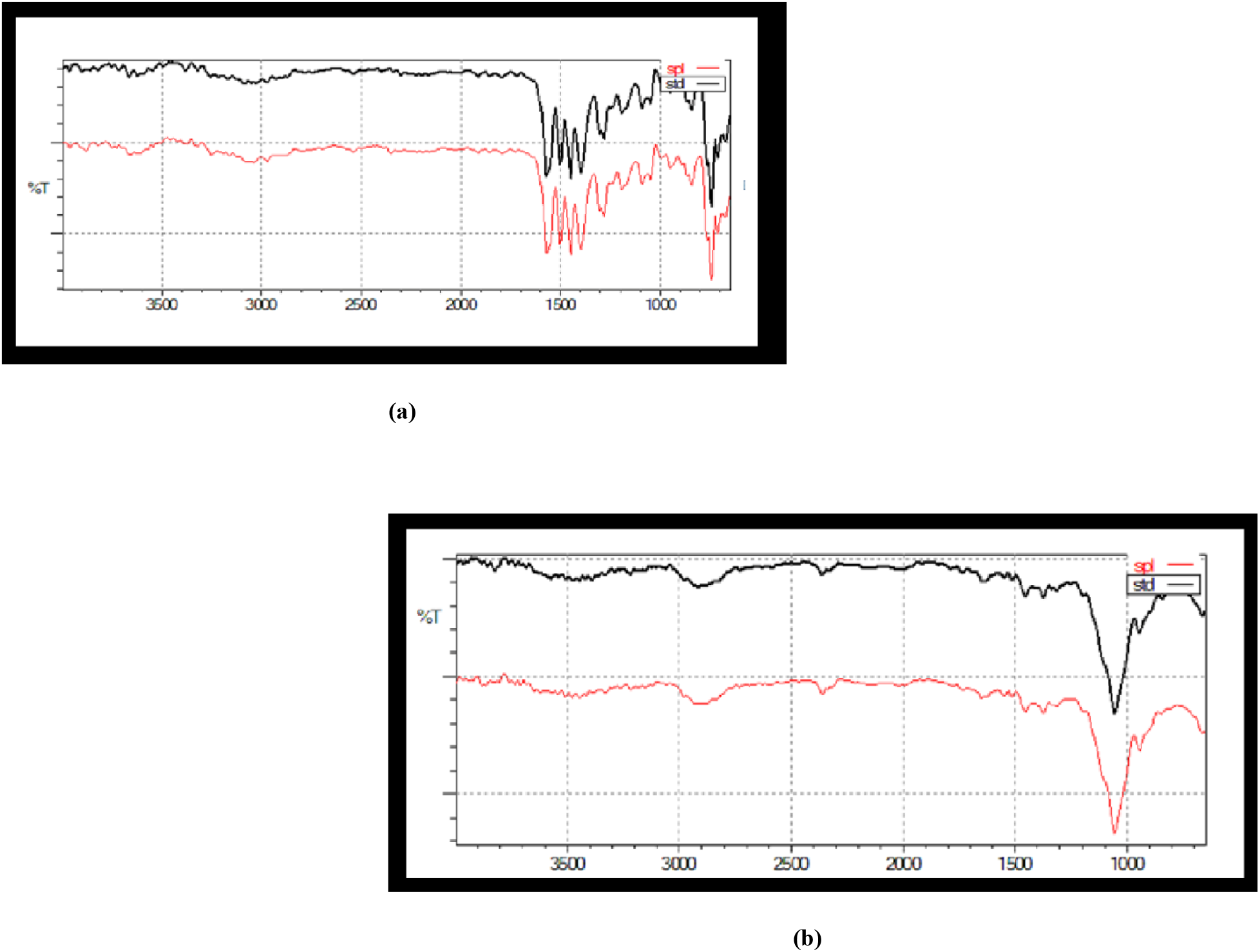
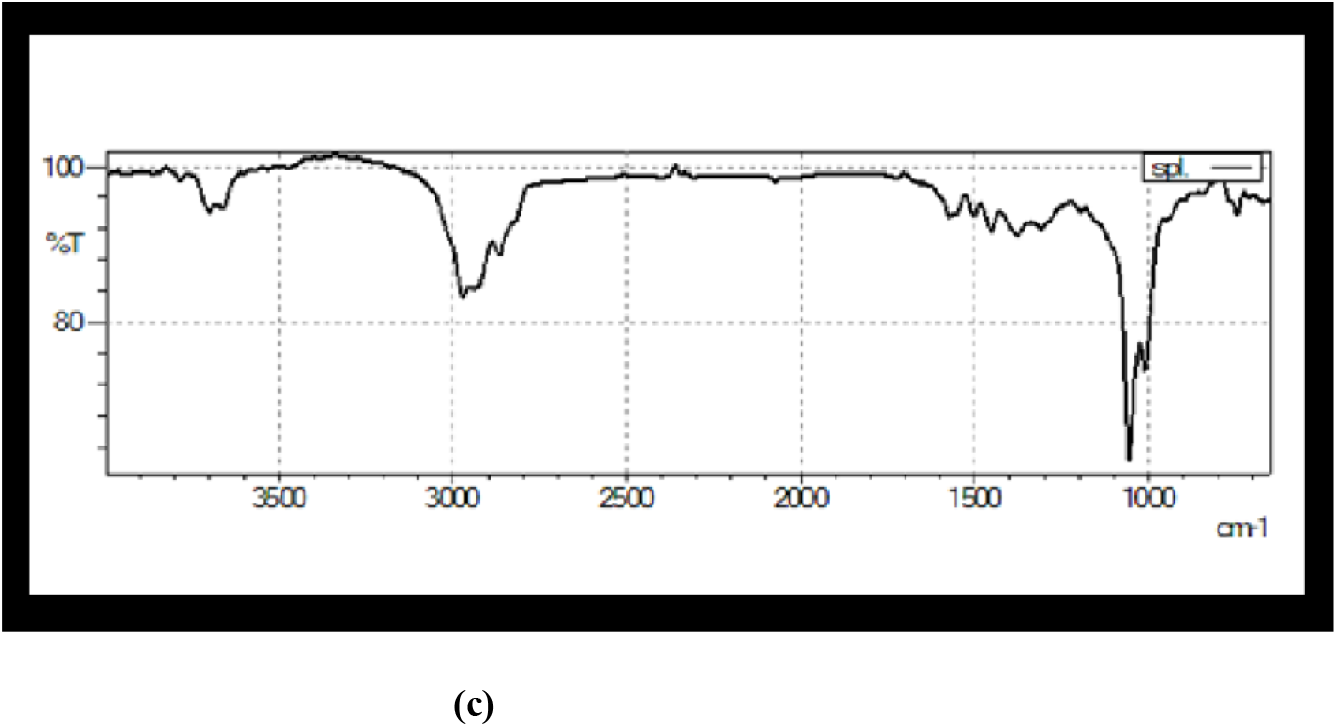
FTIR spectrum of (a) diclofenac standard and sample (b) misoprostol standard and sample (c) diclofenac sodium and misoprostol optimized formulation.

#### 2.3.3 Method of Preparation of compression coated tablets

##### 2.3.3.1 Development of inner core diclofenac tablet

Diclofenac sodium core formulations were designed using two independent variables Sodium starch glycolate (X_1_= 2.5-3.5%) and Povidone K30 (X_2_=5.5-8%) through Design Expert^®^ (7.0.0; Stat-Ease, Inc., USA). The amount of diclofenac sodium (75mg/tablet) and other excipients was kept constant (Table 1). The central composite design was applied as response surface methodology to choose the optimized formulation. Nine formulations of diclofenac sodium were developed; comprising of four factorials, four axial and one center point. The 3D plots were generated to observe the effect of independent variables on tablet hardness and disintegration time. Diclofenac Sodium was mixed with Lactose monohydrate, Corn Starch and with half of the quantity of Sodium Starch Glycolate for 10 minutes by tumbling method. The granulating fluid was prepared by using Povidone K-30 and Isopropyl Alcohol (1:10 ratio); until clear solution was formed. The active powder blend and granulation fluid was mixed in Planetary Mixer (Kenwood, UK) for 3-5 minutes to achieve a damp mass. The damp mass was passed through sieve No. 6 and transferred to the tray dryer for drying at 40ºC. Prepared granules were then passed through the Frewitt oscillating granulator MG-636 fitted with 1.0 mm sieve along with sodium starch glycolate (remaining 50% amount) and magnesium stearate and mixed for 10 minutes and then compress to an average of 145mg ± 7.5% in a local compression machine (Express30).

**Table 1:** Central composite design of Diclofenac sodium (75mg) core tablets and Misoprostol outer tablets (24mg)

| Batch Code | Rotation Points | Coded Levels of variables |  |  |  | Amount of formulation Ingredients (mg) |  |  |  |  |  | Amount per tablet (mg) |
| --- | --- | --- | --- | --- | --- | --- | --- | --- | --- | --- | --- | --- |
|  |  | Sodium starch glycolate | PVP-30 | Sodium starch (%) | PVP-30 (%) | Sodium starch glycolate (mg) | PVP- 30 (mg) | Corn starch | Lactose monohydrate | Magnesium Stearate | Diclofenac sodium |  |
| DF1 | Axial Formulation Run Points | 1 | 1 | 3.5 | 8 | 5.07 | 11.6 | 16.99 | 35.99 | 2.9 | 75 | 147.55 |
| DF2 |  | 0 | 1.414214 | 3 | 8.51777 | 4.35 | 12.35 |  |  |  |  | 147.58 |
| DF3 |  | 1.414214 | 0 | 3.707107 | 6.75 | 5.36 | 9.78 |  |  |  |  | 146.02 |
| DF4 |  | -1 | 1 | 2.5 | 8 | 3.62 | 11.6 |  |  |  |  | 146.1 |
| DF5 | Factorial Formulation Run Points | 0 | -1.41421 | 3 | 4.98223 | 4.35 | 7.22 |  |  |  |  | 142.45 |
| DF6 |  | 1 | -1 | 3.5 | 5.5 | 5.07 | 7.97 |  |  |  |  | 143.92 |
| DF7 |  | -1 | -1 | 2.5 | 5.5 | 3.62 | 7.97 |  |  |  |  | 142.47 |
| DF8 |  | -1.41421 | 0 | 2.292893 | 6.75 | 3.32 | 9.78 |  |  |  |  | 143.98 |
| DF9 | Central Formulation Run Point | 0 | 0 | 3 | 6.75 | 4.35 | 9.78 |  |  |  |  | 145.01 |
| Batch Code | Rotation Points | Coded Levels of variables |  |  |  | Amount of formulation Ingredients (mg) |  |  |  |  |  | Amount per tablet (mg) |
|  |  | Kollidon CL | MCC | Kollidon CL | MCC | Kollidon CL | MCC | Talc | Aerosil | Magnesium Stearate | Misoprostol (1%hpmc dispersion) | Amount per tablet (mg) |
| MF1 | Axial Formulation Run Points | -1 | -1 | 3 | 65 | 14.16 | 306.8 | 1.18 | 0.991 | 1.2272 | 24 | 348.358 |
| MF2 |  | 0 | 0 | 4 | 75 | 18.88 | 354 |  |  |  |  | 400.278 |
| MF3 |  | 0 | -1.41421 | 4 | 60.8579 | 18.88 | 287.212 |  |  |  |  | 333.398 |
| MF4 |  | -1 | 1 | 3 | 85 | 14.16 | 401.2 |  |  |  |  | 442.758 |
| MF5 | Factorial Formulation Run Points | 1.414214 | 0 | 5.414214 | 75 | 25.488 | 354 |  |  |  |  | 406.886 |
| MF6 |  | 1 | 1 | 5 | 85 | 23.6 | 401.2 |  |  |  |  | 452.198 |
| MF7 |  | 0 | 1.414214 | 4 | 89.1421 | 18.88 | 420.75 |  |  |  |  | 467.028 |
| MF8 |  | -1.41421 | 0 | 2.585786 | 75 | 11.8 | 354 |  |  |  |  | 393.198 |
| MF9 | Central Formulation Run Point | 1 | -1 | 5 | 65 | 23.6 | 306.8 |  |  |  |  | 357.798 |

##### 2.3.3.2 Enteric coating of optimized inner core diclofenac tablet

Enteric coating of diclofenac sodium was performed using Eudragit L 100-55, Isopropyl Alcohol and Opadry II Blue 85 F205034. While Polyethylene glycol 6000 was used for polishing of the final tablets. During coating the air inflow temperature was set at 50-55°C and the bed temperature was set at 40° C. The spray volume was kept 50-80g/min. During coating theoretical weight gain of the tablets was 10% ± 2. Some precautions were strictly followed to ensure even coating of the tablets such as tablets should never become very wet during coating rather only remain moist while applying coating solution and air inlet and outlet flaps were kept open constantly. The nozzle head should be checked regularly for any deposition and cleaned with dry brush. The coating dispersion should be stirred continuously for an even application.

##### 2.3.3.3 Development of outer core of immediate release Misoprostol formulations

Misoprostol outer formulations were also designed by Design Expert^®^ (7.0.0; Stat-Ease, Inc., USA) to yield the compression coated tablets. Central composite design was applied to choose the optimized formulation. The composition of all designed formulations is given in table 1. Total nine formulations of misoprostol were developed having 4 factorial, 4 axial and 1 center point. Kollidon CL (X_1_= 3-5%) and microcrystalline cellulose PH 102 (X_2_= 65-85%) were selected as independent variable. The quantity of misoprostol (1% HPMC dispersion) and other excipients was kept constant. To observe the effect of selected excipients on tablet hardness and disintegration time; 3D plots were generated. For tablet manufacturing Misoprostol (1.0% HPMC dispersion), Kollidon CL, Micro crystalline cellulose PH102 and Aerosil-200 were passed through oscillating granulator fitted with 0.8 × 0.5 mm mesh followed by magnesium stearate and talc. The prepared granules and lubricant was then charged in planetary mixer (Kenwood, UK) for 10 minutes.

##### 2.3.3.4 Development of compression coated tablets

The core tablets of misoprostol were compressed containing the optimized formulations of diclofenac sodium (DF9-MF6 and DF9-MF7). The compressional analysis was carried out at 67.38 -107.35MPa to determine the optimal compression pressure at which a stable formulation could be generated. Locally manufactured compression machine fitted with pressure gauge was used for compression. Keeping the tablet weight within ±5% and tablet dimensions constant the effect of different compression pressure was evaluated on yield pressure (Py).

#### 2.3.4 Pharmaceutical Quality Evaluation

Each developed formulation was evaluated on top loading balance (Snow Rex, Taiwan) to check weight uniformity. Other physical parameters including tablet thickness, diameter, hardness, friability were performed as per official standard[18].

##### 2.3.4.1 Disintegration test

Disintegration test for was carried out according to BP specifications using the disintegration apparatus (Erweka ZT-2, Heuesnstanm, Germany) at 37ºC. The time consumed by each immediate release tablet to disintegrate was recorded. The disintegration time for enteric coated diclofenac sodium tablets (n=6) was analyzed in 0.1 N hydrochloric acid, at 37°C (± 1°C), without disks for 120 minutes. Tablets were analyzed for any abrasion, breaking or disintegration before exposing to the alkaline medium. Then same sample along with a plastic disk was operated in disintegration apparatus containing 900ml of buffer pH 6.8 maintained at 37°C (± 1°C) until all tablets in tester disintegrated completely.

##### 2.3.4.2 Assay method

Randomly collected 20 tablets were used for analysis and crushed in mortar and pestle. Solutions of sample and reference standard of Diclofenac sodium were prepared in 0.1 N sodium hydroxide solution. UV Spectrophotometer UV-1800 (Schimadzu, Japan), at 276nm was used to measure the absorbance of sample and reference.

Contents of Misoprostol in tablet formulations were analyzed by liquid chromatographic method. This method employed the use of a pump (LC-10AS, Shimadzu), UV detector (SPDA-10A, Shimadzu) and Lichrospher 100 column (250×4.0 mm, RP-8, 5μm) for quantification of misoprostol at wavelength 210 nm. Mobile phase was prepared with Acetonitrile: IPA: Water (375: 100: 525) and filtered through filter having a porosity of 0.45µm and degas. Standard solution was prepared by taking 500 mg of misoprostol (1% dispersion) working standard and dissolved in 50ml of mobile phase in a 100 ml volumetric flask and the volume was made up. Sample was prepared by taking the upper Misoprostol mantles of randomly selected 50 core tablets. The quantity of mantles equivalent to 25 tablets (5mg Misoprostol) was transferred accurately into a 200 ml of volumetric flask along with 100 ml mobile phase. This mixture was shaken and sonicated for 15 min. Afterward the sample solution was filtered through a filter paper having a porosity of 0.45µm and degassed. Injection volume of standard and sample solution were 5µl. The chromatogram of standard misoprostol is shown in figure 2.

**Figure 2:**
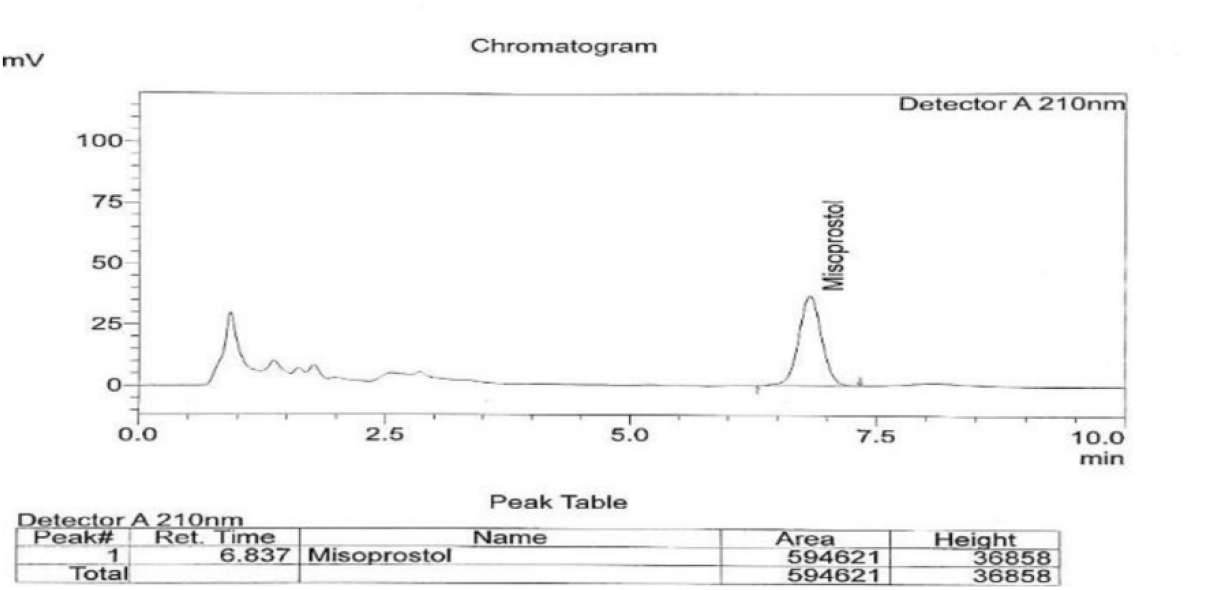
Chromatogram of standard misoprostol.

##### 2.3.4.3 Dissolution test

The dissolution test for enteric coated diclofenac sodium and misoprostol was carried out as per official requirement [18]. The dissolution test of enteric coated diclofenac sodium tablets was performed using rotating Paddle apparatus II (SOTAX AT 7). Initially 6 tablets were placed in 0.1NHCI, maintained at 37±0.5 °C and operated for 2 hours at speed 75 rpm. After 120 minutes, 20 ml of the sample was pipette out for analyzing the drug release by spectrophotometer. The dissolution medium was replaced with phosphate buffer pH 6.8 and same sample of tablets was operated for 1 hour. Aliquot layer of the samples was analyzed to determine drug release spectrophotometrically at 276 nm using phosphate buffer pH 6.8 as blank solution. Percentage drug release of Misoprostol reference (MF6 and MF7) and enteric coated reference tablets of Diclofenac Sodium and optimized DF9 formulation in 0.1 N HCl (pH 1.2) and phosphate buffer (pH 6.8) are shown in figure 3.

**Figure 3:**
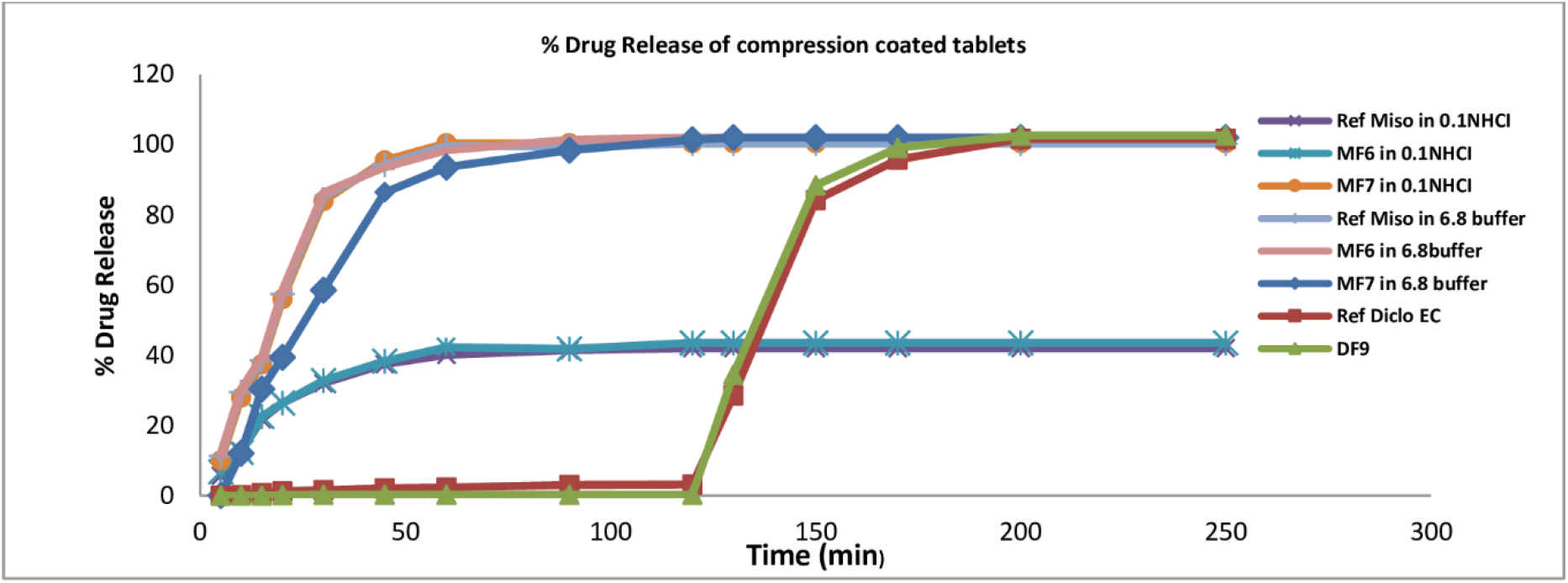
**% Drug release of Misoprostol reference, MF6 and MF7 in 0.1N HCI (pH 1.2) and in phosphate buffer (pH 6.8). % Drug release of enteric coated Reference tablets of Diclofenac Sodium and optimized DF9 formulation in 0.1 N HCl (pH 1.2) and phosphate buffer (pH 6.8)**.

#### 2.3.5 Film coating of misoprostol tablets

Misoprostol tablet formulation MF6 and MF7 were subjected to film coating after compression using Opadry white, Isopropyl Alcohol and Polyethylene glycol 6000. A conventional Coating Pan with Spray gun (local made) was used for coating. The coating parameters were set as follows: batch size 50 g (approx.), air inflow temperature 75-80°C, tablet bed temperature 43– 45°C, pan speed = 40 r/min, spray volume 80-100gm/min (Table 4). When 1/4^th^ portion of coating solution was left Polyethylene glycol 6000 was dissolved in distilled water for polishing purpose to obtain final coated tablets. The total weight gain after coating was 3-5%. Same precautions were followed during coating like tablets should never become very wet, but only remain moist, nozzle head should be checked regularly for any deposition of dried coating solution, coating solutions were stir continuously for even application etc.

#### 2.3.6 Stability studies

Selected tablet formulations were subjected to stability study as per ICH guidelines[19]. Samples were kept in stability chamber at 40±2°C and 75±5% RH and the samples were examined at 0, 3, and 6 months’ interval. The appearance of tablets, disintegration time, drug dissolution and assay were conducted on samples to determine their physical and chemical stability during storage as per pharmacopoeial specification. The data was analyzed for estimating the shelf life using R-Gui software^®^.

## 3. Results and discussion

Gastric damage associated with the use of NSAIDs (e.g. diclofenac sodium) promoted the enteric coating of solid dosage forms. Combination of diclofenac sodium and misoprostol has been recommended to treat pain and inflammation. The aim of this study was to design an effective dosage form for the treatment of rheumatoid arthritis for patients suffering from gastric ulcer. Combining two APIs in a single dosage unit is the way of providing ease of drug administration to the patients suffering from medical complications. Therefore, formulation design has always been challenging for the industrial pharmacist in order to bring innovation in formulation design, ease and convenience for patients.

### 3.1 Micromeritic and FTIR analysis

Micromeritic properties of diclofenac sodium formulations were found as per specification except for DF3 and DF6. The powder blends were evaluated for true density, bulk density, and tapped density and results were found in the range of 1.26-1.40, 0.60-0.63, and 0.55-0.65gm/cm^3^ respectively. For misoprostol formulations the results were found in the range of 1.31-1.43, 0.55-0.61 and 0.53-0.68 gm/cm^3^ respectively. The flowability of the powder blends was also evaluated for inner core and outer core tablets through Hausner’s ratio and angle of repose and their results for diclofenac sodium were lying in the range of 1.16 -1.33 and 31.04 - 44.34 ° respectively. Formulations DF3 and F6 were found only passable and their poor flowability, disqualified them for compression. For outer core formulation of Misoprostol, the results of Hausner’s ratio and angle of repose were found in the range of 1.120-1.36 and 33.23-47.31 ° respectively (Table 2).

**Table 2:** Flowability properties of diclofenac sodium and misoprostol formulations.

| **Batch Code** | **Angle of Repose (n=3)** | **Hausner's Ratio (n=3)** | **Carr's Index (n=3)** | **USP Remarks** |
| --- | --- | --- | --- | --- |
| **DF3** | 44.34±0.264 | 1.33±0.008 | 24.54±0.877 | Passable |
| **DF6** | 43.35±0.575 | 1.31±0.0182 | 23.87±0.636 | Passable |
| **DF7** | 32.3±0.448 | 1.16±0.040 | 14.73±0.636 | good |
| **DF8** | 31.04±0.665 | 1.150±0.012 | 11.73±0.773 | good |
| **DF9** | 32.86±0.503 | 1.117±0.055 | 15.00±0.784 | good |
| **Batch Code** | **Angle of Repose (n=3)** | **Hausner's Ratio (n=3)** | **Carr's Index (n=3)** | **USP Remarks** |
| **MF2** | 47.31±0.350 | 1.36±0.012 | 27.732±0.773 | Poor |
| **MF4** | 37.75±0.585 | 1.123±0.055 | 19.245±0.784 | Fair |
| **MF5** | 46.66±0.503 | 1.37±0.018 | 26.49±1.105 | Poor |
| **MF6** | 33.28±0.665 | 1.120±0.008 | 11.43±0.877 | Good |
| **MF7** | 33.23±0.514 | 1.142±0.040 | 12.46±0.636 | Good |

Formulations which did not yield acceptable results of micromeritic properties or which were exceeding the weight limit form the target formulation (DF1, DF2, DF4, DF5, and MF2, MF4, and MF5) were also not selected for further study. FTIR analysis of dicolofenac sodium standard and its sample, misoprostol standard and sample and optimized formulation of miso-diclo is shown in the Figure 1 (a, b and c respectively). The spectrum of optimized formulation exhibited an intense, well defined peak and the infrared band nearly 1602.79 cm^-1^(C=C) and 3209.62 cm^-1^(COOH) peak and 3387.00 cm^-1^(N-H amine) peaks were reported. As per the IR spectra of optimized formulation interpreted that there was no shifting in the frequencies of stated functional groups. Thus on the basis of above given results it is concluded that no there was no significant drug and excipients interaction was found.

### 3.2 Pharmaceutical Quality Attributes

#### 3.2.1 Formulation design and characterization

Diclofenac sodium tablets were designed by Design of Experiment^®^ software and manufactured by wet granulation. These tablets showed good flow ability, uniform drug distribution, low weight variation, and good mechanical strength. All physical parameters of core tablets were complied with the official standard. Average weight of core tablets was maintained at 145.0mg ± 7.5%, thickness 3.02-3.06mm, hardness 9.98-19.9Kg; friability 0.25-0.72 %, disintegration time 7:11 - 16:11minutes, drug release 79.48-97.17% and assay 82.34-101.36% (Table 3). Tablets with increased hardness and disintegration time were excluded from further investigation and stability testing was performed on DF7, DF8 and DF9 formulations.

**Table 3:** Quality attributes of Diclofenac sodium core and Misoprostol formulation.

| Batch Cod<br>(core tabs) | Weight<br>Variation<br>(mg)<br>Mean ± SD<br>(n=20) | Thickness<br>(mm)<br>Mean ± SD<br>(n=20) | Hardness<br>(Kg)<br>Mean ± SD<br>(n=20) | Friability<br>(%) | Disintegration<br>time<br>(min) | Drug Release<br>(%)<br>Mean ± SD<br>(n=6) | Assay<br>(Mean ± SD<br>(n=3) |
| --- | --- | --- | --- | --- | --- | --- | --- |
| DF3 | 146.52± 2.86 | 3.050±0.047 | 19.98±1.136 | 0.25 | 16.11 | 81.66 ± 2.16 | 83.74± 1.57 |
| DF6 | 143.14± 1.79 | 3.022±0.063 | 18.99±1.183 | 0.34 | 15.33 | 79.48 ± 4.22 | 82.34±1.86 |
| DF7 | 142.88 ± 2.92 | 3.057±0.053 | 9.98±1.239 | 0.72 | 11.23 | 97.17±1.57 | 101.36±1.07 |
| DF8 | 143.17± 1.59 | 3.041±0.059 | 10.32±1.430 | 0.47 | 8.1 | 92.45±1.35 | 99.61±1.60 |
| DF9 | 145.76± 2.64 | 3.063±0.052 | 12.27±1.293 | 0.39 | 7.11 | 96.72±1.48 | 101.33±1.35 |
| Batch Code<br>(core tabs) | Weight<br>Variation<br>(mg)<br>Mean ± SD<br>(n=20) | Thickness<br>(mm)<br>Mean ± SD<br>(n=20) | Hardness<br>(Kg)<br>Mean ± SD<br>(n=20) | Friability<br>(%) | Disintegration<br>time<br>(min) | Drug Release<br>(%) in<br>phosphate<br>buffer<br>Mean ± SD<br>(n=6) | Assay<br>(Mean ± SD<br>(n=3) |
| MF4 | 450.01 ± 1.92 | 5.681 ± 0.002 | 14.45±1.162 | 0.65 | 16.01 | 84.560±2.316 | 88.66±1.527 |
| MF6 | 465.88 ± 3.18 | 5.890 ± 0.003 | 11.45±1.239 | 0.41 | 14.88 | 95.617±1.587 | 99.166±1.040 |
| MF7 | 470.77 ± 2.92 | 5.945 ± 0.02 | 12.88±1.430 | 0.49 | 12.11 | 101.445±1.025 | 102.60±1.630 |
| Compression<br>coated<br>formulation | Weight<br>Variation<br>(mg)<br>Mean ± SD<br>(n=20) | Thickness<br>(mm)<br>Mean ± SD<br>(n=20) | Hardness<br>(Kg)<br>Mean ± SD<br>(n=20) | Friability<br>(%) | Disintegration<br>time<br>(min) | Drug Release<br>(%) in<br>phosphate<br>buffer<br>Mean ± SD<br>(n=6) | Assay<br>(Mean ± SD<br>(n=3) |
| DF9-MF7 | 490.53± 1.65 | 6.111 ± 0.008 | 13.88 ± 2.14 | 0.21 | 13.43 | 101.25 ± .143 | 100.83 ± 1.22 |

Formulation DF9 exhibited satisfactory results during stability studies with shelf life 4.8years using R-Gui software ^®^. Therefore, DF9 was selected for enteric coating in the next step of this study. The inherent mechanical strength of core tablets selected for enteric coating should be sufficiently high so that during coating process any defect such as abrasion, erosion, or crumbling may not affect the overall integrity of the tablets and shows inherent weakness [20,21]. The coating process was performed in conventional coating pan employing spray coating technique. There are certain important factors for ensuring the smoothness and uniformity of coated tablets such as temperature of coating pan and spray volume [22]. These process parameters can impact the film formation during coating and may influence the disintegration and dissolution test results [23,24]. The coating procedure for enteric coating yielded NMT 10% of weight gain (Table 4). The average weight of batches was maintained at 160mg ± 7.5 %. The effectiveness of enteric coating was evaluated by disintegration and dissolution test. Disintegration time of coated tablets was examined as per USP procedure and none tablet showed any abrasion or disintegration in acidic medium for 2hour; but in phosphate buffer of pH 6.8 the disintegration was found in the range of 21:00-22:40minutes [25]. The drug dissolution of DF9 formulation in 0.1NHCI after 120 minutes was found less than 10%, and in phosphate buffer pH 6.8; the drug dissolution after 60 minutes was found in the range of 84.56-101.445% as per official criteria (Figure 3) [22]. The assay result of all batches was found in the range of 88.66-102.60% (Table 3).

**Table 4:**
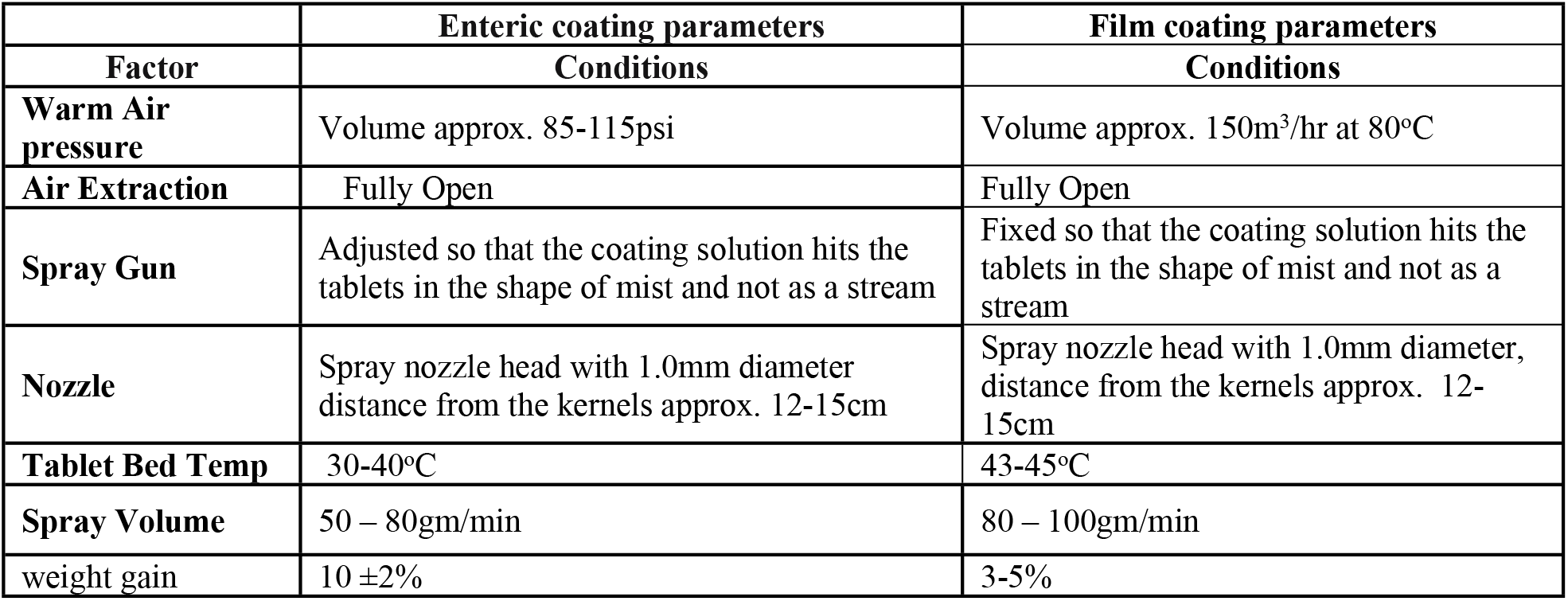
Parameters for enteric coating and film coating.

|  | **Enteric coating parameters** | **Film coating parameters** |
| --- | --- | --- |
| **Factor** | **Conditions** | **Conditions** |
| **Warm Air pressure** | Volume approx. 85-115psi | Volume approx. 150m<sup>3</sup>/hr at 80°C |
| **Air Extraction** | Fully Open | Fully Open |
| **Spray Gun** | Adjusted so that the coating solution hits the tablets in the shape of mist and not as a stream | Fixed so that the coating solution hits the tablets in the shape of mist and not as a stream |
| **Nozzle** | Spray nozzle head with 1.0mm diameter distance from the kernels approx. 12-15cm | Spray nozzle head with 1.0mm diameter, distance from the kernels approx. 12-15cm |
| **Tablet Bed Temp** | 30-40°C | 43-45°C |
| **Spray Volume** | 50 – 80gm/min | 80 – 100gm/min |
| **weight gain** | 10 ±2% | 3-5% |

These batches were stored in stability chamber for stress stability testing over period of six months. Satisfactory results were obtained showing no substantial variation in physical and chemical testing of the enteric coated tablets. Thus result indicates that employed manufacturing process is consistent. Based on the satisfactory physicochemical results the next step was covering of formulations DF9 with Misoprostol (1% HPMC dispersion) to develop final formulation as compression coated tablet.

#### 3.2.2 Formulation design of outer tablets of Misoprostol

In the second phase of this study, outer core tablets of Misoprostol (1.0% HPMC dispersion) were designed by Design of Experiment^®^ using Kollidon CL, Avicel PH 102, Talc, Aerosil-200 and Magnesium Stearate and final tablets were manufactured by wet granulation. The outer mentle of Misoprostol was designed at average weight of 472mg/tablet. This mentle was then used to cover the optimized enteric coated diclofenac sodium tablets to yield compression coated tablet. Three formulations were selected to formulate the final compression coated tablets based on good micromeritic properties and sufficient weight. Formulations with very low amount of powder blend (MF1, MF3, MF8, and MF9) were excluded form study as they were unable to cover the inner enteric coated tablets. Formulations MF2 and MF5 showed poor results of micromeritics, and therefore these were not further included in study. Pharmaceutical quality attributes of all compression coated formulations (final) were found as per official standards. All formulation passed the assay test. The tablet hardness was found to be 11.45-14.45kg, % friability: 0.41-0.65% however, MF4 failed to comply with compendial requirements of % drug release (84.56%) (Table 3). Formulation MF6 and MF7 (Figure 3) showed better results of drug release and were subjected to stability testing. Formulations bearing code MF6 and MF7 were processed for enclosing the enteric coated diclofenac sodium tablets to yield the compression coated tablets.

The optimized formulations were subjected to film coating using Opadry white and Polyethylene glycol 6000. Results of these formulations after film coating (final compression coated tablets) have been mentioned in table 3. Amongst all DF9-MF7 formulation was designated as compression coated tablet with satisfactory quality attributes and shelf life of 4.8 years. Furthermore, this formulation did not show any discoloration during accelerated storage conditions and found as the best formulation.

#### 3.2.3 Stability Studies

Stability studies of misoprostol tablets were assessed as per ICH guidelines. Three replicates of optimized formulation were stored at 40 ±2°C and 75 ± 5% RH in the humidity chamber for six months [26]. Following the storage period, the samples were analyzed for drug content, disintegration time and in vitro dissolution rate. After stability testing on these formulations the data was statistically analyzed through R-Gui software^®^ for evaluating the shelf life of final dosage form.

#### 3.2.4 Yield pressure determination

Yield pressure (Py) of optimized formulation was evaluated by reported method at different compression pressure by using Heckel equation [27,28]. It has been accepted that materials showing Py value less than 80 are soft or plastic and those having Py value more than 80 are hard or brittle [29]. However, this classification for each material type depends upon the variation in conducted experiments and other factors; yet no standardized method is available to characterize the material under study. As Heckel plots are not ideally linear so another factor which contributes variation in reported Py value is the region from where this value is derived. In the present study for formulation MF6 and MF7 the values of Py were found to be 833.33 and 909.09 MPa indicating brittle nature of compacts (Figure 4).

**Figure 4:**
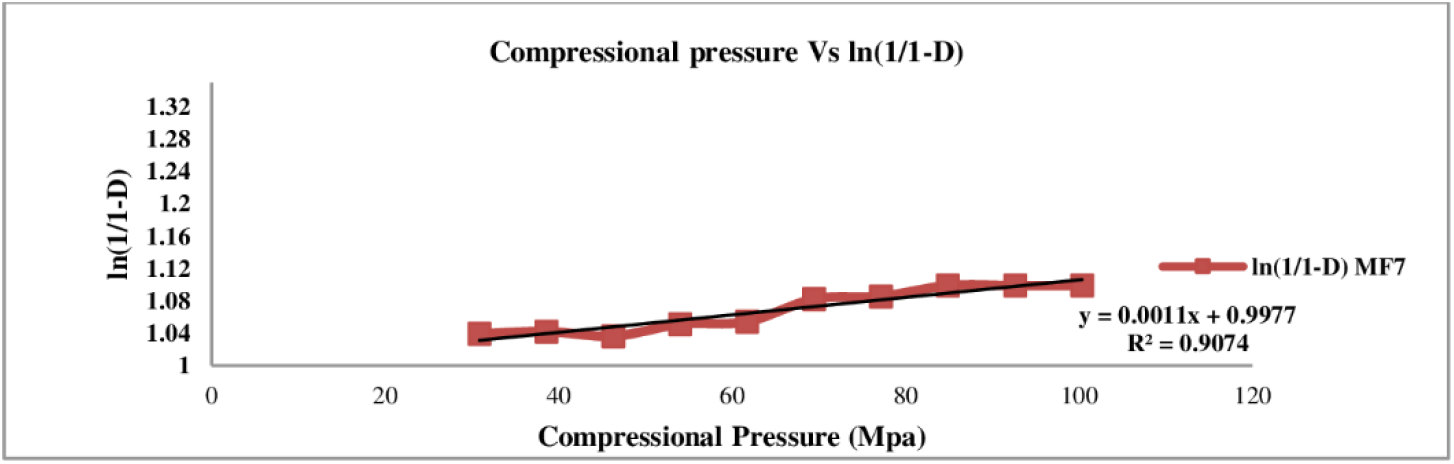
Compressional analysis of optimized compression coated formulation.

A relationship between applied compressional pressure and tablet hardness was established showing that at pressure 69.48-77.20MPa the tablet hardness was found satisfactory (Figure 5).

**Figure 5:**
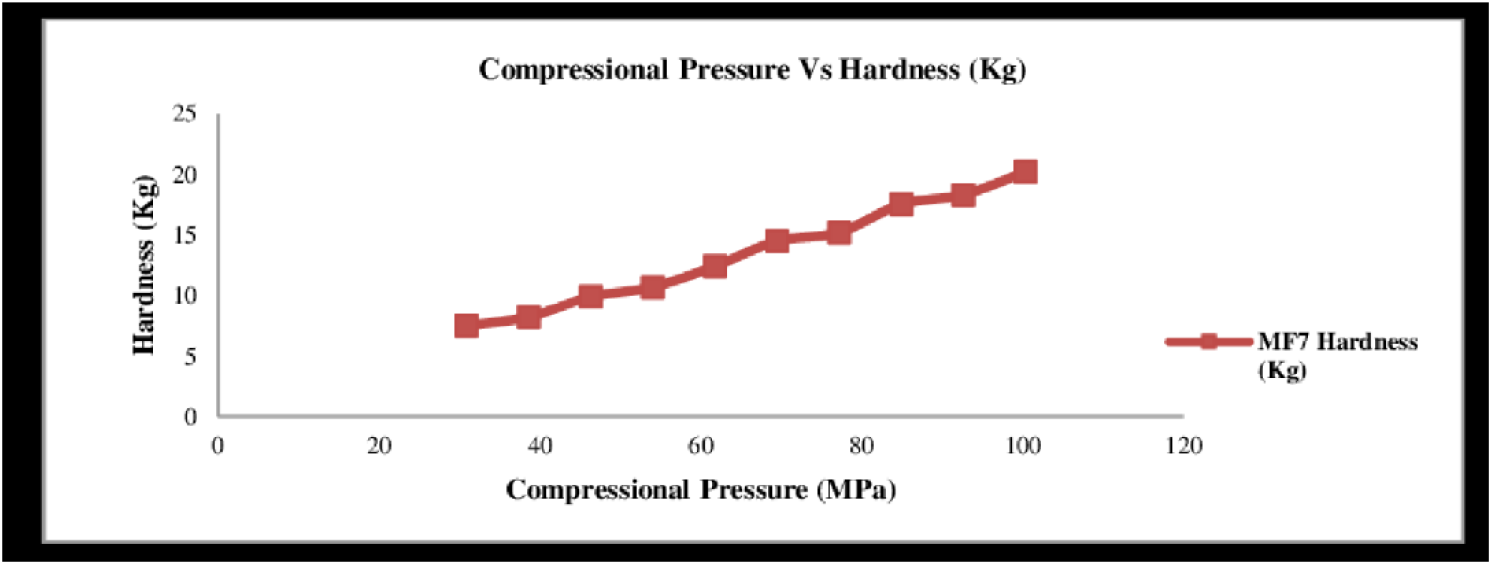
Applied compression pressure Vs Hardness of compression coated tablets.

Beyond this compressional pressure an increased in tablet hardness was observed along with prolonged disintegration time. Therefore, it is suggested to keep the applied compressional pressure within 4-5tons (i.e. 69.48-77.20 MPa) to produce the tablets of sufficient mechanical strength.

#### 3.2.5 RSM Plots

The interaction effect of independent variable on the responses was observed from RSM plots obtained using design expert^®^ 7.0.0 software and illustrated in figure 6 and 7 for diclofenac sodium and misoprostol respectively. The response relation i.e. disintegration with independent variables (sodium starch glycolate and PVP K30) is showing negligible response between PVP K30 content and disintegration time while sodium starch glycolate showed significant reduction in disintegration time as the amount of sodium starch was increased in the formulations (Figure 6a). In Figure 6b, it can be clearly depicted that as the proportion of PVP K-30 is increased in the formulation, tablet hardness was also increased. This effect is observed due to the fact that it has good binding property. Although, there was no significant response observed on addition of the sodium starch glycolate.

**Figure 6:**
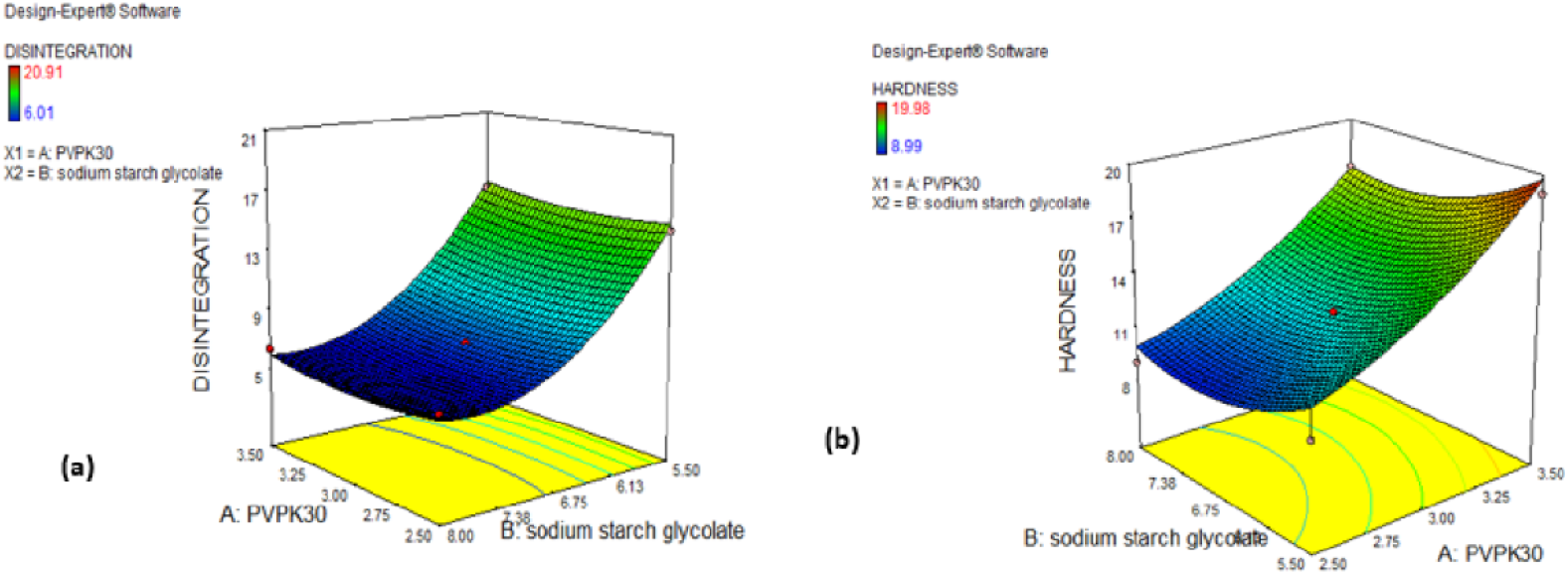
3D Response surface plots of diclofenac formulations presenting effect of independent variables on (a) disintegration time, (b) hardness.

**Figure 7:**
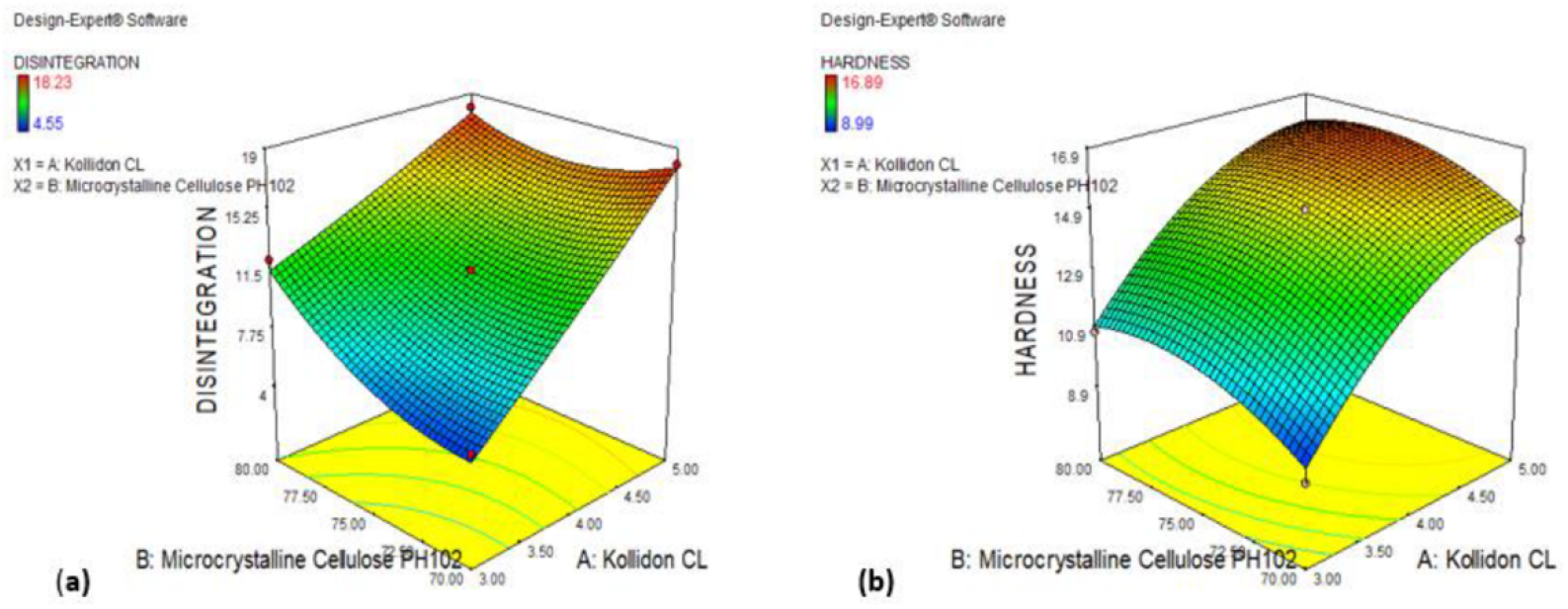
3D Response surface plots of misoprostol formulations presenting effect of independent variables (a) disintegration time, (b) tablet hardness.

For misoprostol the response relation i.e. disintegration with independent variables (Kollidon CL and Avicel) is mentioned in figure 6a. Figure 6b explains that the presence of Kollidon CL increased the tablet hardness as its proportion was increased in the formulation. Furthermore, the formulation was prepared through wet granulation method and hardness was found within the recommended standard specification. However, avicel did not contribute significantly to the tablet hardness. Figure 7a explains that the presence of Kolliodon CL increases the disintegration time with increase in its concentration while avicel slightly contributed to improvement in disintegration time. In nut shell wet granulation and carefully selected excipients contributed in improving the mechanical strength of inner tablet.

## 4. Conclusion

Compression coated tablets of Diclofenac sodium and Misoprostol have been successfully developed by wet granulation and optimized by Design of Experiment. Designed formulations showed satisfactory quality attributes. Amongst all formulation DF9-MF7 was chosen as optimized formulation based on satisfactory quality attributes and compressional analysis. This formulation can also be subjected to in-vivo study in future on human volunteers for good management of arthritis.

## Notes

### Competing Interest Statement

The authors have declared no competing interest.

